# Elevated somatic mutation burdens in normal human cells due to defective DNA polymerases

**DOI:** 10.1101/2020.06.23.167668

**Authors:** Philip S. Robinson, Tim H.H. Coorens, Claire Palles, Emily Mitchell, Federico Abascal, Sigurgeir Olafsson, Bernard Lee, Andrew R.J. Lawson, Henry Lee-Six, Luiza Moore, Mathijs A. Sanders, James Hewinson, Lynn Martin, Claudia M.A. Pinna, Sara Galvotti, Peter J. Campbell, Iñigo Martincorena, Ian Tomlinson, Michael R. Stratton

**Affiliations:** Wellcome Sanger Institute, Hinxton, CB10 1SA, UK; Department of Paediatrics, University of Cambridge, Cambridge, CB2 0QQ, UK; Institute of Cancer and Genomic Sciences, University of Birmingham, B15 2TT, UK; Department of Pathology, Queen Mary Hospital, Hong Kong University, Hong Kong; Department of Haematology, Erasmus University Medical Centre, 3015 CN, Rotterdam, The Netherlands; Edinburgh Cancer Research Centre, IGMM, University of Edinburgh, Crewe Road, Edinburgh EH4 2XZ, UK

**Author notes:** These authors contributed equally (Philip S. Robinson, Tim H.H. Coorens, Claire Palles). These authors jointly supervised this project (Ian Tomlinson, Michael R. Stratton).

## Abstract

Mutation accumulation over time in normal somatic cells contributes to cancer development and is proposed as a cause of ageing. DNA polymerases Pol ε and Pol δ replicate DNA with high fidelity during normal cell divisions. However, in some cancers defective proofreading due to acquired mutations in the exonuclease domains of POLE or POLD1 causes markedly elevated somatic mutation burdens with distinctive mutational signatures. POLE and POLD1 exonuclease domain mutations also cause familial cancer predisposition when inherited through the germline. Here, we sequenced normal tissue DNA from individuals with germline POLE or POLD1 exonuclease domain mutations. Increased mutation burdens with characteristic mutational signatures were found to varying extents in all normal adult somatic cell types examined, during early embryogenesis and in sperm. Mutation burdens were further markedly elevated in neoplasms from these individuals. Thus human physiology is able to tolerate ubiquitously elevated mutation burdens. Indeed, with the exception of early onset cancer, individuals with germline POLE and POLD1 exonuclease domain mutations are not reported to show abnormal phenotypic features, including those of premature ageing. The results, therefore, do not support a simple model in which all features of ageing are attributable to widespread cell malfunction directly resulting from somatic mutation burdens accrued during life.

## INTRODUCTION

Replication of the genome is required at each cell division. It is effected by DNA polymerases synthesising a new DNA strand with a sequence dictated by a template strand. Low error rates are ensured by the fidelity of base incorporation, proofreading capabilities of the polymerases and surveillance by the DNA mismatch repair machinery. DNA replication in humans is largely performed by the polymerases Pol ε and Pol δ which undertake leading and lagging strand synthesis respectively^1,2^.

Uniquely among nuclear polymerases, both Pol ε and Pol δ have proofreading activities, mediated by their exonuclease domains, which identify and remove mismatched bases^1,3-5^. Somatically acquired heterozygous missense mutations in the POLE or POLD1 exonuclease domains found in some human cancers cause defective proofreading and, consequently, high burdens of somatic mutations with distinctive mutational signatures^6-9^. Cancers with POLE exonuclease domain mutations show very high single base substitution (SBS) mutation burdens whereas cancers with POLD1 exonuclease domain mutations show less elevated SBS burdens but are often associated with microsatellite instability^8^. Mutations generated by defective proofreading POLE and POLD1 show marked replication strand bias consistent with their differential roles in leading and lagging strand synthesis^1-2,8^. Polymerases with these mutations also cause mutator phenotypes when engineered into yeast and mice^10-15^.

POLE and POLD1 exonuclease domain mutations can also be inherited through the germline causing a rare autosomal dominant familial cancer predisposition syndrome, known as Polymerase Proofreading Associated Polyposis (PPAP), characterised primarily by early-onset colorectal and endometrial tumours^16,17^. It is plausible that an increased somatic mutation rate underlies this cancer predisposition and high somatic mutation loads have been reported in the small number of neoplasms analysed from such individuals^17^. However, whether the mutation rate is elevated in normal cells, or just in neoplastic cells, is not known. If elevated in normal cells, the magnitude of the increase, whether it is raised over the whole lifespan, the range of tissues and fraction of cells in each tissue it affects, and the impact of subsequent neoplastic change are important questions to address in elucidating the pathogenesis of neoplastic transformation.

Accrual of somatic mutations has been proposed as the primary biological mechanism underlying ageing^18-21^. This hypothesis is based on the premises that a) mutations accumulate throughout life; and b) higher mutation loads cause widespread malfunction of cell biology. Recent reports have confirmed that the somatic mutation burden in normal cells does increase during life in a more or less linear manner^22-30^, compatible with a causal role for somatic mutations in ageing. However, somatic mutations could, in principle, accumulate without significant biological consequences. Thus, study of individuals with inherited POLE or POLD1 exonuclease domain mutations could provide insight into the wider biological consequences of elevated mutation burdens and the pathogenesis of ageing.

## RESULTS

### Clinical information and samples

Fourteen individuals, aged between 17 and 72 years, each carrying one of four different germline exonuclease domain mutations in POLE or POLD1 (POLE L424V (n=8), POLD1 S478N (n=4), POLD1 L474P (n=1) and POLD1 D316N (n=1)), were studied. Eleven had a history of five or more colorectal adenomas, with age at first polyp diagnosis ranging from 15 to 58 years. Five were diagnosed with colorectal cancer, all before age 50, and all had a known family history of colorectal adenoma, colorectal cancer and/or other cancers. No other consistent phenotypic abnormalities were reported^17^ (Extended Data Table 1).

### Mutagenesis in normal intestinal stem cells

The epithelial cell population of an intestinal crypt is a clone derived from a single ancestral crypt stem cell that existed <10 years prior to sampling^31-34^. Somatic mutations found in the large majority of cells in a crypt, and thus with a high variant allele fraction (VAF), recapitulate the set of mutations present in that ancestral cell^26^. Thus, to investigate somatic mutation burdens and rates in normal cells from POLE and POLD1 germline mutation carriers, 109 normal intestinal crypts (colorectum n=85, ileum n=10 and duodenum n=14) were individually isolated by laser capture microdissection from biopsy and surgical resection samples of 13 individuals and whole-genome sequenced (WGS) (median 33.5-fold coverage) (Methods, Extended Data Table 2).

The somatic SBS burdens in the seven individuals with POLE L474V and four with POLD1 S478N correlated with age, indicating that mutation accumulation is likely to be continuous through life, linear and at similar rates in each individual carrying the same mutation. Crypts from individuals with POLE L424V showed an average SBS mutation rate of 331/year (linear mixed-effects model 95% confidence interval (C.I.) 259-403, *P*= 10^−12^) (**Fig. 1a and 1b**, Methods, Supplementary code). The POLD1 S478N germline mutations were associated with an SBS rate of 152/year (linear mixed-effects model 95% C.I. 128-176, *P*= 10^−17^) and POLD1 D316N and L474P were associated with an SBS rate of 58/year (linear mixed-effects model 95% C.I. 51-65, *P*= 10^−22^). By comparison, intestinal crypts from normal individuals acquire 49 SBS per year^26^ (linear mixed-effects model 95% C.I. 46-52, *P*= 10^−36^) (Methods). Therefore increased somatic SBS rates are present in all normal intestinal cells of individuals with POLE or POLD1 germline mutations (**Fig. 1**), although there are differences in mutation rates between POLE (∼7-fold higher than normal individuals) and POLD1 (up to 3-fold higher) germline mutations, and between different POLD1 mutations. Indeed, individuals with POLD1 D316N and L474P exhibited relatively modest elevations of SBS rates (∼1.2-fold). There was also evidence of differences in SBS rates between the five individuals with POLE L424V suggesting the existence of genetic and/or environmental modifiers of mutation rate (**Fig**.**1b**, Methods, Supplementary Code).

**Figure 1.**
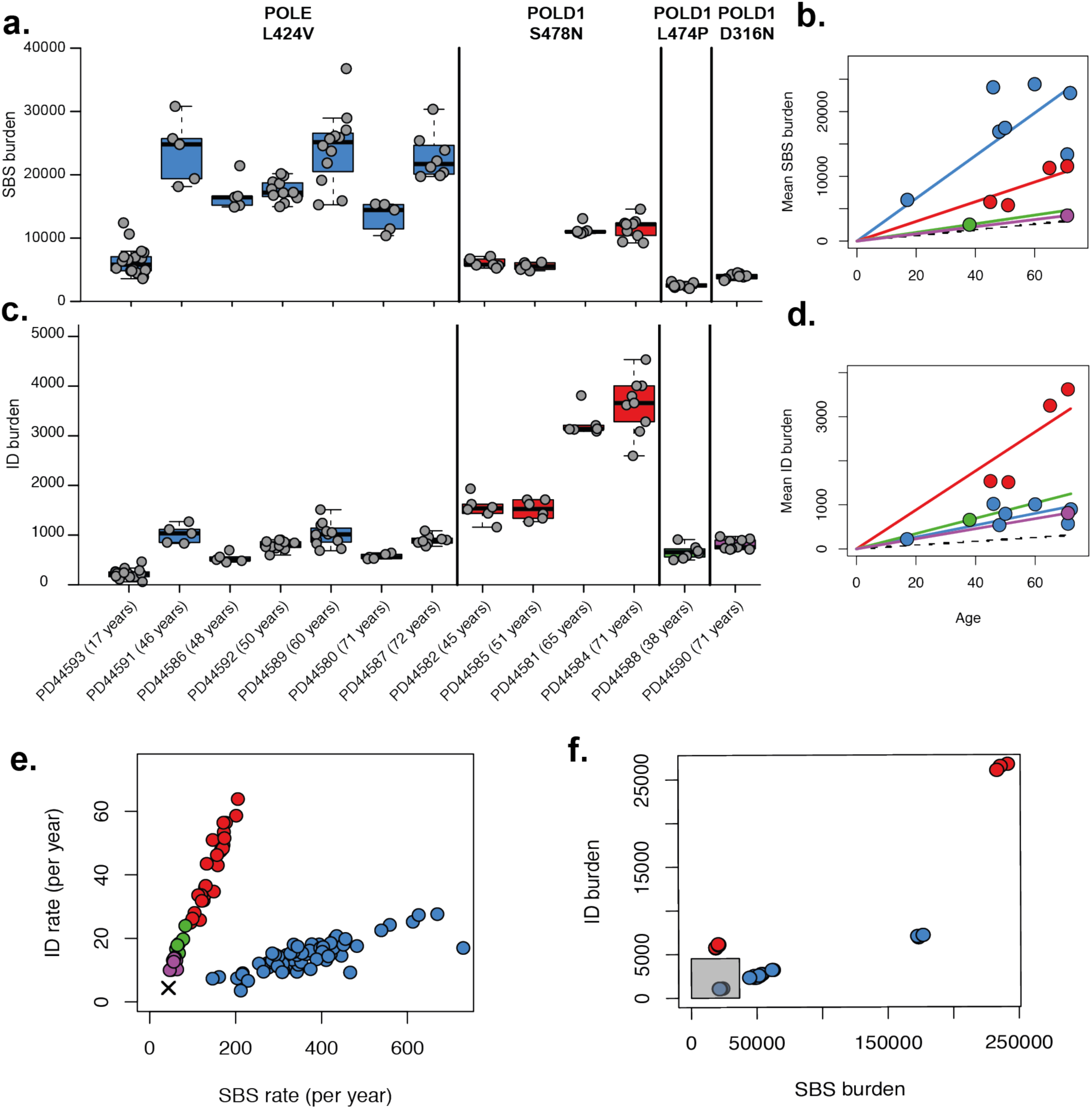
SBS and ID burdens in normal and neoplastic intestinal crypts from individuals with germline *POLE* or *POLD1* mutations. **(a)** Genome-wide mutation burden per individual, with the specific germline mutation indicated on top and colour-coded (blue, red, green and purple for POLE L424V, POLD1 S478N, POLD1 L474P, and POLD1 D316N, respectively. Boxplots display median, inter-quartile range (IQR) from 1^st^ to 3^rd^ quartiles and whiskers extend from the last quartile to the last data point that is within 1.5x IQR. **(b)** Mean SBS burden versus age, showing regression lines for the four different germline mutations. The relationship between age and SBS burden in normal individuals is displayed as a dashed line. **(c)** The genome-wide ID burden per individual. **(d)** The relationship between age and ID burden. **(e)** ID rate (per year) versus SBS rate (per year), the cross marks the SBS and ID rate in normal individuals. **(f)** ID and SBS burden in adenomatous samples from individuals with POLE/POLD1 mutations. The grey box indicates the range of mutation burdens in normal intestinal crypts from individuals with POLE/POLD1 mutations.

Small insertion and deletion (ID) mutation rates in normal intestinal crypts were also elevated in individuals with germline POLE/POLD1 mutations, with rates of 13/year (POLE L424V), 44/year (POLD1 S478N) and 12/year (POLD1 D316N and POLD1 L474P) (linear mixed-effects model 95% C.I., 10-16, 35-53, 9-16, *P*=10^−10^,*P*=10^−13^ and *P*=10^−9^ respectively). These are all substantially above the expected rate of 1/year in individuals without POLE or POLD1 mutations^26^ (**Fig. 1c and 1d**) (Methods, Supplementary Code). Individuals with POLE and POLD1 germline mutations therefore showed differences in their relative rates of accumulation of base substitutions and indels (**Fig. 1e**). Normal cells from individuals with POLE mutations exhibited 28-fold higher SBS than ID mutation rates, whereas cells from individuals with POLD1 mutations showed 4-fold higher rates, consistent with previous findings in cancer genomes and experimental systems^6,8,11,35-37^. With the exception of one individual, who had been treated with oxaliplatin for colorectal cancer, and one with an incidental finding of trisomy of the X chromosome (47 XXX) (Extended Data Fig. 1a and Fig.2), copy number changes and rearrangements were rare, occurring at a similar prevalence to normal crypts from normal individuals. Reductions in telomere length with age also occurred at similar rates to normal crypts from normal individuals (Extended Data Table 2, Methods, Supplementary Code).

Crypt-like structures from six colorectal adenomas and one carcinoma from individuals with germline POLE or POLD1 mutations were also microdissected and sequenced. SBS and ID burdens were considerably higher than in normal crypts from the same individuals sampled at the same time (**Fig. 1e and 1f**), albeit with substantial variation between lesions. Therefore, increases in SBS and ID mutation rates are associated with the conversion from a normal to an adenoma or cancer crypt in individuals with POLE or POLD1 germline mutations, a similar pattern to that observed in normal individuals^38,39^.

### Mutational signatures

Eleven SBS mutational signatures were observed in normal intestinal crypts from individuals with POLE and POLD1 germline mutations (**Fig. 2a-c**, Extended Data Fig. 3). Nine have been previously reported; SBS1, SBS5, SBS10a, SBS10b, SBS17b, SBS28, SBS35, SBSA and SBSB. SBS1 (characterised by C>T substitutions at N<u>C</u>G trinucleotides and likely due to deamination of 5-methylcytosine) and SBS5 (of unknown etiology) are found in all normal intestinal crypts from normal individuals where they accumulate in a more-or-less linear manner with age^7,9,26,40^. SBSA and SBSB are found in some normal intestinal crypts from some normal individuals and are predominantly acquired during childhood^26,41^. SBSA is likely due to colibactin, a mutagenic product of a strain of *E*.*coli* sometimes present in the colon microbiome^26,42^. SBS10a, SBS10b and SBS28 were previously found in the subsets of colorectal, endometrial and other cancer types with somatically acquired POLE mutations^7,9^ (**Fig. 2a**). Two therapy-associated signatures were identified; an SBS35-like signature which is associated with platinum-based chemotherapy^9,43^ in an individual treated with oxaliplatin (**Fig. 2c**, Extended Data Fig. 1b-e) and a signature characterised predominantly by T>G mutations in an individual treated with capecitabine^43,44^(**Fig. 2f**). Two new mutational signatures (SBS10c and SBS10d) were observed in normal and neoplastic crypts from individuals with germline POLD1 mutations. Both were characterised predominantly by C>A substitutions; in SBS10c at A<u>C</u>C, C<u>C</u>A, C<u>C</u>T, T<u>C</u>A and T<u>C</u>T trinucleotides and in SBS10d at T<u>C</u>A and T<u>C</u>T trinucleotides (the mutated base is underlined) (**Fig. 2d and 2e**, extended sequence context shown in Extended Data Fig.4a).

**Figure 2.**
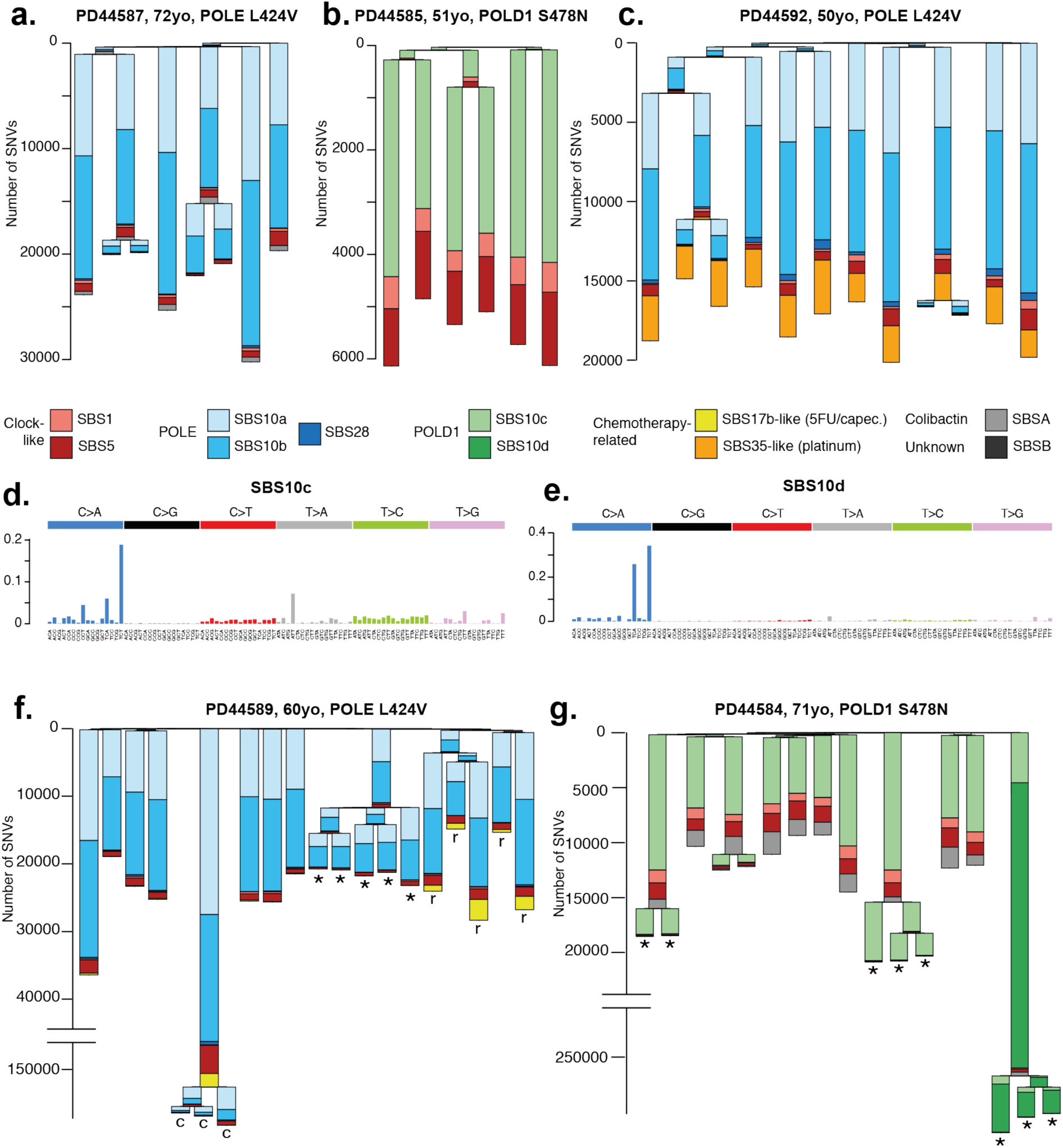
Phylogenies of intestinal crypts with mutational signature annotation. Phylogenies of microdissected intestinal crypts of **(a)** PD44587 (POLE L424V), exhibiting mainly SBS10a and SBS10b, and **(b)** PD44585 (POLD1 S478N), exhibiting SBS10c. SBS1 and SBS5, normal signatures of ageing, are also present. Signature exposures are colour-coded as indicated below the trees. Branch lengths correspond to SBS mutation burdens. **(c)** In addition to mutagenesis due to POLE L424V, PD45592 shows widespread exposure to mutagenesis due to a platinum-based chemotherapeutic agent (SBS35-like). Probability distributions of SBS10c **(d)** and SBS10d **(e)**, two novel signatures associated with POLD1 mutagenesis. **(f)** Phylogeny of PD44589 (POLE L424V) containing samples from driver-bearing adenomas (marked with ‘*’) and a carcinoma (marked with ‘c’). Note that the y-axis is broken for scale. This individual showed mutagenesis due to exposure to capecitabine (SBS17b-like), which was localised to carcinoma samples and nearby normal rectum (marked with ‘r’). **(g)** Phylogeny of PD44584 (POLD1 S478N), with driver-bearing adenomas marked with ‘*’. One particular polyp showed extensive hypermutation (note broken y-axis), largely due to SBS10d.

The increases in somatic base substitution burdens in normal intestinal crypts from POLE germline mutation carriers compared to normal individuals were almost completely attributable to SBS10a, SBS10b and SBS28 mutations and those in POLD1 carriers to SBS10c mutations. By contrast, the estimated burdens of SBS1, SBS5, SBSA and SBSB found in normal intestinal crypts from *POLE*/*POLD1* germline mutation carriers were similar to those expected in normal individuals of the same ages. Thus, defective POLE/POLD1 proofreading appears not to substantially affect the rates of the mutational processes underlying SBS1, SBS5, SBSA and SBSB. POLE and POLD1 are responsible for leading and lagging strand DNA synthesis respectively^1,2^. Consistent with these roles, there was marked replication strand bias of SBS10a and SBS10b somatic mutations in POLE germline mutation carriers with the opposite bias of SBS10c and SBS10d in POLD1 mutation carriers (Extended Data Fig. 4b-f).

The elevated mutation loads in adenoma and carcinoma crypts from POLE/POLD1 germline mutation carriers compared to normal intestinal crypts from each individual were also predominantly due to increased burdens of SBS10a and 10b (in POLE mutant cases) and SBS10c and SBS10d (in POLD1 mutant cases) (**Fig. 2f and 2g**). However, crypts from POLE polyps showed greater relative increases in SBS10a than SBS10b (**Fig. 2a and 2b**). The mechanisms underlying these marked accelerations in mutation rates during neoplastic change, and why they apply differentially to the processes underlying the different signatures, are unknown.

IDs in normal intestinal crypts from both POLE and POLD1 germline mutation carriers were dominated by single T insertions at T homopolymer tracts, characteristic of signature ID1. ID1 mutations were also further increased in neoplastic crypts compared to normal crypts from each individual (**Figure 1c and 1f**, Extended Data Fig.5, Extended Data Table 2).

Cancer “driver” mutations (Methods) found in crypts from normal intestine and colorectal neoplasms from individuals with POLE/POLD1 germline mutations showed similar SBS and ID mutational spectra to genome-wide spectra from normal intestinal crypts from these individuals (Extended Data Fig. 6a-c, Extended Data Table 3). The proportion of normal crypts with drivers in POLE and POLD1 mutation carriers was not different from those in unaffected individuals (Extended Data Fig.6d).

### Mutagenesis in other tissues

Cancers of the colorectum and endometrium are the predominant types associated with germline and somatic POLE/POLD1 mutations^16-17,45^. Similar to intestinal crypts, endometrial glands are clones derived from a single recent ancestral stem cell and whole genome sequencing of a gland reveals the mutations present in that ancestral cell^29,46^. Twelve endometrial glands dissected from a 60 year-old carrying a POLE L424V germline mutation showed elevated rates of SBS accumulation (177/year vs. 29/year in normal individuals) and ID (13/year vs. <1/year in normal individuals) (**Fig 3a**). SBS10a and SBS10b were responsible for the increase in SBS rate compared to normal individuals (**Fig 3b**). Somatic driver mutations in cancer genes are common in normal human endometrium^25-29,30^ and were found in a similar repertoire of genes in all but one gland from this individual (Extended Data Fig. 6e, Extended Data Table 3).

**Figure 3.**
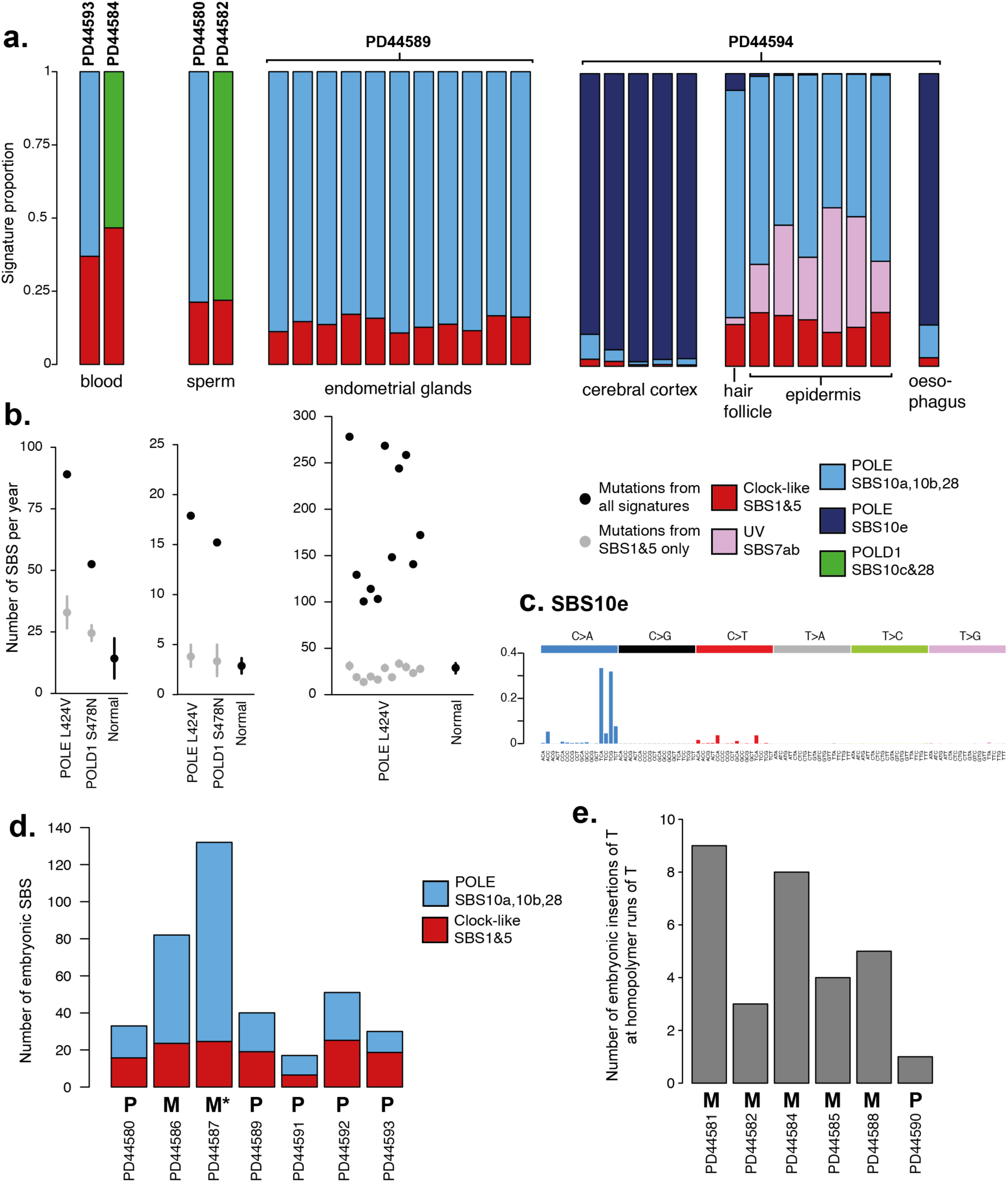
POLE and POLD1 mutagenesis in other tissues. **(a)** Signature contribution to mutational landscapes of various tissues in individuals with a POLE L424V (PD44593, PD44580, PD44589, PD44594) or POLD1 S478N (PD44584, PD44582) germline mutation. Normal blood and sperm were sequenced using a modified duplex sequencing protocol, while other tissues were subjected to low-input WGS after laser-capture microdissection. Groups of mutational signatures are colour-coded as indicated. **(b)** Estimated genome-wide total mutation rate per year for blood, sperm, and endometrium (black dots), as well as yearly mutation burden due to SBS1 and SBS5 (grey dots with 95% confidence intervals). Normal mutation rates for blood^48^ and endometrium^29^ are displayed for reference, as well as the paternal age effect for *de novo* germline mutations as a proxy for the mutation rate of sperm^47^. **(c)** Probability distribution for SBS10e, a novel signature identified in PD44594 and associated with POLE mutagenesis in a subset of tissues. **(d)** Early embryonic SBSs in individuals with a POLE L424V germline mutation, with contribution from POLE signatures (blue, SBS10a, SBS10b, SBS28) and normal signatures (red, SBS1 and SBS5). ‘M’ indicates the mutation was inherited maternally, ‘P’ paternally. ‘M*’ indicates presumed maternal inheritance based on pedigree. **(e)** Early embryonic insertions of T at homopolymers of T (indicative of POLD1 mutagenesis) in individuals with POLD1 germline mutations (S478N: PD44581, PD44582, PD44584, PD44585; L474P: PD44588; D316N: PD44590). Again, ‘P’ and ‘M’ indicate paternal and maternal inheritance, respectively.

We also microdissected and sequenced fragments of cerebral cortex, skin epidermis and oesophageal squamous epithelium from a single 46 year old individual with germline POLE L424V (PD44594). The VAFs of mutations from these samples indicate that they contain multiple cell clones and thus mutation burdens are difficult to directly estimate (Extended Data Table 2). Nevertheless, mutagenesis due to defective POLE proofreading was found in all tissues (**Fig 3a**). In skin, ageing-and UV-related signatures were accompanied by SBS10a and SBS10b. In both brain and oesophagus, a further novel mutational signature (SBS10e) was present, characterised predominantly by C>A substitutions at T<u>C</u>G and T<u>C</u>A trinucleotides (**Fig 3c**). The mechanism underlying this tissue-limited signature is not known and this signature has not previously been observed in extensive surveys of human cancer and several normal tissue types^7,9,23,26-29,41^.

To further investigate mutagenesis in other cell types and embryonic germ layers, we used a modified duplex sequencing protocol on DNA from whole blood and sperm. By sequencing single DNA molecules at low error rates this method allows quantification of mutation burdens from tissues in which cells derived from many progenitors are intimately mixed and clonal units for sequencing cannot be dissected (Methods). The results from four individuals, two with POLE L424V and two with POLD1 mutations, showed elevated SBS rates in blood and sperm compared to normal controls. Estimated yearly mutation rates due to SBS1 and SBS5 in blood and sperm were consistent with previous estimates^47,48^ and the excess mutation burdens were accounted for by SBS10a and SBS10b (POLE mutant individuals) and SBS10c (POLD1 mutant individuals) (**Fig 3b**). The data suggest that mutagenesis due to defective POLE/POLD1 is additive to normal ageing.

In summary, mutational signatures associated with POLE/POLD1 exonuclease domain mutations were found in all cells from all tissues examined and across the three germ layers. The elevation of burden was variable between tissues, being higher in intestinal crypts and endometrial glands than in other tissues. The results from sperm indicate that the elevated mutation rate extends beyond somatic tissues into the germline.

### Mutagenesis during early embryogenesis

Somatic mutations accumulate throughout development, from the first cell division onwards^49-52^. A mutation arising in an early embryonic cell may be present in a substantial proportion of adult cells and in multiple different tissues^49^. Early embryonic mutations can be detected in whole-genome sequences of highly polyclonal adult tissue samples as mutations with relatively high VAF^49-50,52^. Using this approach, putative early embryonic mutations were identified from whole-genome sequences of whole-blood samples. The embryonic mutational spectra of some POLE mutant individuals exhibited large exposures to SBS10a, SBS10b and SBS28, whereas others were dominated by SBS1 and SBS5, the signatures normally causing early embryonic mutations^52^ (**Fig 3d**). Similarly, in POLD1 mutant cases, the number of early embryonic single base pair insertions was highly elevated in some, but not all individuals (**Fig 3e**). This heterogeneity reflects the inheritance pattern of the germline mutation and is likely a consequence of the maternal to zygotic transition of gene expression^53^. When a germline POLE/POLD1 mutation is paternally inherited, any effect on mutagenesis is delayed until zygotic genome activation, thus sparing the early embryo for the first few cell divisions. When maternally inherited, however, the defective proofreading polymerase is present in the cytoplasm of the ovum and POLE/POLD1 mutagenesis therefore occurs immediately after fertilisation, leading to a high prevalence of such mutations in early embryogenesis. These results indicate that mutagenesis due to defective POLE/POLD1 proofreading is present at the earliest stages of life.

### Differential mutation burdens across the genome

We compared the distribution of somatic mutations across the genome in individuals with germline POLE/POLD1 mutations to those found in unaffected individuals. To achieve this, we used the inherent SBS1 and SBS5 exposure as a baseline for normal ageing associated processes and calculated the genome-wide fold increase in mutations due to the additive effects of POLD1 and POLE mutagenesis (**Fig 4a**). Mutation burdens due to the various forms of SBS10 were heavily biased towards late replicating regions (Extended Data Fig. 7). Proportionately, this spares protein-coding exons which are mainly located in early replicating regions. This relatively constrained increase in mutation burdens in protein-coding exons may conceivably mitigate the biological consequences of the elevated somatic mutation rates due to POLE/POLD1 germline mutations. Nevertheless, differential burdens between tissues were maintained and mutation rates in coding regions were more increased in colon and endometrium than in skin (**Fig 4b**).

**Figure 4.**
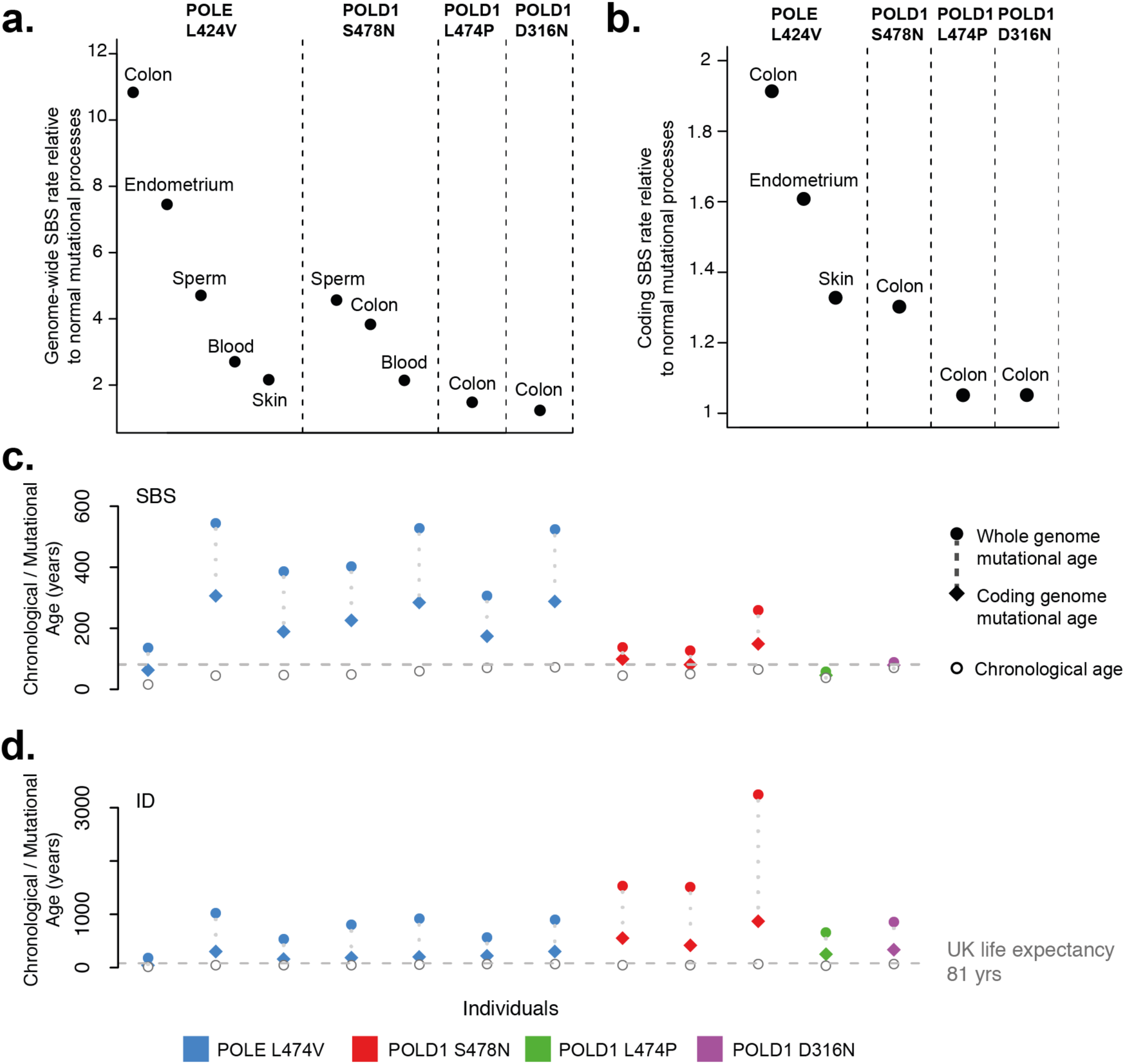
Increase of mutation burdens genome-and exome-wide. **(a)** Fold increase of genome-wide mutation burden due to POLE and POLD1 germline mutations across various normal tissues. Fold increases were calculated as total mutagenesis due to both POLE (SBS10a, SBS10b, SBS28) or POLD1 (SBS10c, SBS28) and normal ageing (SBS1, SBS5, and for skin SBS7a and SBS7b) divided by mutagenesis due to normal ageing only. **(b)** The same calculation restricted to exonic regions only, showing a much lower fold increase than found genome-wide. Mutation burdens in blood and sperm were too low to adequately quantify exonic mutational signature proportions. **(c-d) “**Mutational ages” and chronological ages of histologically normal intestinal crypts displayed per individual. “Mutational ages” are calculated based on the expected rate of mutation accumulation in wild-type intestinal crypts^26^ enabling the calculation of both SBS and ID “mutational ages”. Plots display the SBS **(c)** and ID **(d)** “mutational ages” across the whole genome (filled dots) and coding genome (filled diamonds). Germline mutation is colour-coded; blue, red, green and purple for POLE L424V, POLD1 S478N, POLD1 L474P, and POLD1 D316N, respectively. Individuals’ chronological ages are indicated by unfilled circles and United Kingdom life expectancy is displayed as a dashed horizontal line.

## DISCUSSION

This study shows that multiple normal cell types from POLE/POLD1 exonuclease domain germline mutation carriers demonstrate the mutational signatures and elevated somatic SBS and ID mutation rates characteristic of defective proofreading by these polymerases. The results are consistent with the presence of elevated mutation rates in all cells of all types throughout life.

The extent of the elevation in mutation rate appears greater in intestinal and endometrial epithelium than in the other cell types analysed. The basis for this variation is not understood, but may reflect different stem cell division rates. It may also, at least in part, explain the predilection for colorectal and endometrial cancer observed in these individuals.

The somatic mutation theory of ageing proposes that the increasing somatic mutation burdens in normal cells, continuously accrued over a lifetime, have increasingly detrimental effects on cell function and thus engender the set of phenotypic features collectively termed ageing^18-21^. The mutation burdens observed in cells from POLE/POLD1 mutation carriers are higher than those of normal individuals of the same ages. Therefore, POLE/POLD1 mutation carriers have elevated “mutational ages” (**Fig. 4c-d**). The biological consequences of this generalised elevated mutation burden appear, however, to be relatively limited. Other than the increase in incidence of colorectal, endometrial and other neoplasms, phenotype information from more than 100 POLE/POLD1 mutation carriers does not obviously reveal features of premature ageing and many survive into the late decades of standard human lifespan^16,17^(Extended Data Table 2). Therefore, the rare natural experiment of germline POLE/POLD1 exonuclease domain mutations leading to elevated mutation rates does not support a simple somatic mutation theory of ageing. The results are, moreover, similar to those obtained in mice with engineered germline POLE and POLD1 exonuclease domain mutations^10,11^.

Important cautions, however, should temper this conclusion. First, more comprehensive measurement of somatic mutation burdens across cell types in POLE/POLD1 germline mutation carriers is indicated. The varying degrees of mutation rate elevation between cell types potentially leaves some, which could be particularly influential in generating the ageing phenotype, relatively protected. Second, POLE/POLD1 exonuclease domain germline mutation carriers show similar burdens of somatic copy number changes, rearrangements and telomere erosion to normal individuals (Extended Data Table. 2, Supplementary Code). If ageing depends on these mutation classes, it would not be accelerated in POLE/POLD1 mutation carriers. Finally, additional factors may mitigate the impact of elevated mutation burdens in POLE/POLD1 exonuclease domain carriers. For example, a disproportionately small fraction of the mutation burdens due to SBS10a,b,c,d,e falls in coding regions of the genome, potentially reducing its biological impact^54^.

Nevertheless, the results indicate that many normal human cell types throughout life tolerate high SBS and ID mutation rates and therefore that direct effects of somatic mutation accumulation may not underlie all components of the progressive biological dysfunction termed ageing.

## METHODS

### Human tissue and blood samples

Human tissue and blood was collected under United Kingdom NHS Research Ethics Committee approval (17/SC/0079). Tissue was collected under informed consent from patients during routine endoscopy and from surgical resection samples (PD44580-PD44593) and in one case following autopsy (PD44594). Peripheral blood samples were collected at various time points into ethylenediaminetetraacetic acid (EDTA). All samples were stored at -80 °C.

### DNA extraction from bulk samples

Frozen whole blood underwent DNA extraction using the Gentra Puregene Blood Kit (Qiagen). Briefly, 1-2mls of frozen blood were thawed, lysed in RBC lysis solution and centrifuged. Cell pellet was resuspended in cell lysis solution and incubated at 37 °C for 2 hours. RNA and protein was degraded using RNase A solution and protein precipitation solution. After these stages, protein precipitation solution was added and the sample was agitated and centrifuged. DNA was precipitated with isopropanol. DNA was extracted from semen samples using B-mercaptoethanol followed by phenol chloroform extraction^55^.

### Tissue Preparation

Tissues were embedded in Optimal Cutting Temperature (OCT) compound, frozen histological sections were cut at 30µm and mounted on polyethylene naphthalate (PEN) slides and fixed in 70% ethanol for 5 minutes followed by two washes with phosphate buffered saline for 1 minute each. Slides were manually stained in haematoxylin and eosin using a conventional staining protocol. A subset of samples (PD44594c-h and PD44589f) were fixed in PAXgene Tissue FIX (Qiagen) according to manufacturer’s instructions. Fixed tissue samples were embedded in paraffin using a Tissue-Tek tissue processing machine (Sakura). No formalin was used in the preparation, storage, fixation or processing of samples. Processed tissue blocks were embedded in paraffin wax, sectioned to 10µm thickness and mounted onto PEN slides (Leica). Tissue slides were stained using a standard haematoxylin and eosin (H&E) protocol. Slides were temporarily cover-slipped and scanned on a NanoZoomer S60 Slide Scanner (Hamamatsu), images were viewed with NDP.View2 software (Hamamatsu).

### Laser Capture Microdissection

Laser capture microdissection was undertaken using a LMD7000 microscope (Leica) into a skirted 96-well PCR plate. Cell lysis was undertaken using 20µl proteinase-K Picopure® DNA Extraction kit (Arcturus®), samples were incubated at 65 °C for 3 hours followed by proteinase denaturation at 75 °C for 30 minutes. Thereafter samples were stored at -20 °C prior to DNA library preparation.

### Intestinal crypt isolation

Crypts from one tissue block (PD44593e) were isolated using EDTA chelation. In brief, dissected mucosa was incubated in an EDTA solution and gently agitated resulting in dissociation of intestinal crypts from the underlying components of the intestinal epithelium. Crypts were then separated under a light microscope and placed in ATL buffer (Qiagen) containing 10%(v/v) proteinase K and digested overnight at 56°C. DNA extraction was performed using the QiaAMP DNA micro kit (Qiagen) as per manufacturer’s instructions. DNA was then stored at -20°C.

### Low-input DNA library preparation and sequencing

DNA library preparation of micro-dissected tissue samples was undertaken as previously described using a bespoke low-input enzymatic-fragmentation-based library preparation method^26-27,29^. This method was employed as it allows for high quality DNA library preparation from very low starting quantity of material (from 100-500 cells). DNA library concentration was assessed after library preparation and used to guide choice of samples to take forward to DNA sequencing, minimum library concentration was 5ng/µL and libraries with >15ng/µL were preferentially chosen. 150bp paired-end Illumina reads were prepared with Unique Dual Index barcodes (Illumina).

DNA sequencing was undertaken on a NovaSeq 6000 platform using XP kit (Illumina). Samples were multiplexed in pools of 6-24 samples. Pools were sequenced to achieve a coverage of ≥30x.

### Mutation calling and post-processing filters

Sequencing reads were aligned to NCBI human genome GRCh37 and aligned using the Burrow-Wheeler Alignment (BWA-MEM)^5^. Single Base Substitutions (SBS) were called using the ‘Cancer Variants through Expectation Maximization’ algorithm (CaVEman)^6^. Mutations were called using an unmatched normal synthetic bam file to retain early embryonic and somatic mutations. Post-processing filters were applied to remove low-input library preparation specific artefacts and germline mutations using a previously described method^28-29,52^. Filters applied were: (1) common single nucleotide polymorphisms were removed by filtering against a panel of 75 unmatched normal samples^56^ (2) to remove mapping artefacts a minimum alignment score of reads that support a mutation was applied (ASMD ≥ 140) and fewer than half of the read should be clipped (CLPM =0); (3) a filter to remove overlapping reads that result from the relatively short insert size which could lead to double counting of variant reads; and (4) a filter to remove cruciform DNA structures that can arise during the low-input library preparation method.

Next, mutations were aggregated per patient and a read pile-up was performed using an in-house algorithm (cgpVAF) to tabulate the read count of mutant and reference reads per sample for each mutation locus. Germline mutations were filtered out using an exact binomial test and further artefacts by quantifying the beta-binomial overdispersion parameter (rho) for variants across all samples from the same patient and setting a threshold for rho>0.1 for genuine variants^7,8^. The code for these filters can be found at https://github.com/TimCoorens/Unmatched_NormSeq. Phylogenetic trees were created using MPBoot (version 1.1.0 bootstrapped - 1000) and mutations were mapped to branches using maximum likelihood assignment.

Indels (ID) were called using Pindel^57^ using the same synthetic unmatched normal sample employed in SBS mutation calling. ID calls were filtered to remove calls with a quality score of <300 (‘Qual’; sum of mapping qualities of the supporting reads) and a read depth of less than 15. Thereafter, ID filtering was performed in a similar manner as SBS to remove germline variants and library preparation /sequencing artefacts.

### Copy-number alteration calling

Somatic copy-number variants (CNVs) were called using the Allele-Specific Copy number Analysis of Tumours (ASCAT) algorithm^58^, https://github.com/Crick-CancerGenomics/ascat) in the ascatNGS package^59^. Bulk (blood or in one case tissue) samples were used as matched normals. ASCAT was initially run with default parameters. To reduce the number of false-positive calls that arise in normal tissue samples, a segmentation penalty was applied in the ASCAT ‘aspcf’ step. Optimum performance was observed with a penalty value of 100 which was subsequently applied to all samples. Copy-number calls were further filtered to remove artefacts. Copy-number (CN) calls less than 2MB were excluded. Samples with a goodness-of-fit of less than 95% were excluded. CN calls at specific recurrent breakpoints were removed. Sharing of CNVs between samples from different tissue blocks and across individuals that violated phylogenetic structures implied from SBS and ID phylogenetic trees were treated as artefactual and removed from analysis. Similarly, any recurrent copy-number calls with identical break points that were observed across different individuals were also removed. CNV calls were manually verified by visualisation of reads in JBrowse^60^.

### Structural variant calling

Whole-genome sequences were analysed for somatic structural variants (SVs) using Breakpoints via assembly (BRASS) algorithm^61^, paired blood samples were used as controls. If no blood sample was available, a tissue sample was used that was phylogenetically distant to the sample being analysed. SV calls were filtered using an in-house algorithm in a multi-stage process^29^. Finally, all SV calls were manually inspected to confirm somatic variants. SV calls in L1 transposon donor regions and fragile sites were excluded from the final SV analysis.

### Mutational signature analysis

The R package HDP (https://github.com/nicolaroberts/hdp), based on the hierarchical Dirichlet process^62^, was used to extract mutational signatures. Analysis of mutational signatures using this package has been applied to normal tissues previously^26-29^. In brief, this nonparametric Bayesian method models categorical count data using the hierarchical Dirichlet process. A hierarchical structure is established using patients as the first tier (parent nodes) and individual samples as the second tier (dependent nodes). Uniform Dirichlet priors were applied across all samples. The algorithm creates a mutation catalogue for each sample and infers the distribution of signatures in any one sample using a Gibbs sampler. We performed mutational signatures analysis per-branch, counting each branch of the phylogenetic tree as a distinct sample to avoid double counting of mutations. Since the MCMC process scales linearly with the number of counts, we randomly subsampled each branch to a maximum of 2500 total substitutions. Branches with fewer than 100 mutations were excluded from the mutational signature extraction. No reference signatures were included as priors.

Fourteen signature components were extracted (Supplementary Information). A subset of these 14 signatures appeared to be combinations of previously reported reference signatures^9^. To deconvolute composite signatures and to equate obtained HDP signatures to reference ones, we assessed the cosine similarity and employed an EM-algorithm to deconstruct these signatures into reference constituents (SBS1, SBS5, SBS7a, SBS7b, SBS10a, SBS10b, SBS17a, SBS17b, SBS25, SBS28, SBS31, SBS35,^9^ as well as SBSA and SBSB^26^).

In this way, HDP3 was broken down into SBS1 and SBS5; HDP4 into SBS10a and SBS5, HDP5 into SBS1, SBS5, SBS10a, and SBS28; HDP9 into SBS7a, SBS7b, SBS10a, and SBS28 (Supplementary Information). HDP1 was broken down into SBS10a and SBS10b, but SBS10b poorly reflected the observed C>T component of HDP1 and HDP6, which heavily impacted later fitting of signatures. Hence, we constructed our version of SBS10b by subtracting the estimated contribution of SBS10a from HDP1. In this vein, HDP1 became SBS10a and SBS10b, and HDP6 was decomposed into SBS10b, SBS1, SBS5, and SBS28.

HDP2, 7 and 13, exclusively found in patients with a POLD1 germline mutation, were renamed as SBS10c-e, respectively, and not subjected to decomposition. HDP0, 8 and 11 only had one major contributor and were replaced by the purer reference signatures SBS5, SBSA, and SBSB. HDP10 was found to be due to platinum-based chemotherapy, as it is constituted of SBS31 and SBS35. However, the report of a spectrum of signatures due to these chemotherapy agents^43^ and the relatively poor reconstitution using SBS31 and SBS35 only, prompted us to retain HDP10 without further decomposition hence, we name it SBS35-like. A similar approach was used for the capecitabine-related HDP12 component, which resembles SBS17b but has closer similarity to previously reported therapy-related signatures^43,44^. Hence, we renamed HDP12 SBS17b-like.

Therefore, we identify a total of 14 signatures: SBS1, SBS5, SBS7ab, SBS10a-e, SBS17b-like, SBS28, SBS35-like, SBSA, and SBSB. These signatures were refitted to all mutation counts of branches of phylogenies using the R package sigfit (https://github.com/kgori/sigfit)^63^. To avoid overfitting, a limited subset of reference mutational signatures were included per patient corresponding to the HDP signatures that have been identified in that individual. In the case of SBS10d, it was only fitted to branches in which an exposure had originally been reported.

### Validation and extension of mutational signature analysis

In addition to HDP, we also used the non-negative matrix factorisation (NMF) based algorithm SigProfiler^9^ to extract SBS mutational signatures. SigProfiler reports many fewer substitution signature components (7 vs 14) (Supplementary Information), but the components it does extract have clearly recognisable counterparts in the compendium of HDP components (Supplementary Information). Additional components that are stably extracted and have close resemblance to known, reported signatures were identified by HDP but not SigProfiler.

To assess the association between signatures in different mutational classes and to validate the findings of previous iterations of signature extraction, combined extraction of SBS, ID and doublet based substitutions (DBS) was performed using *de novo* extraction with HDP, leading to a direct incorporation of ID and DBS components in SBS10ab, SBS10c, SBS7ab, SBS35-like and SBSA (Supplementary Information).

### Mutational signature assignment

To assess the likely causative mutational signatures for individual mutations, all SBS mutations that mapped branches of the phylogenetic trees were assigned signature probabilities. These assigned probabilities were used to subset mutations for further analysis i.e. replication strand and extended sequence context biases as displayed in Extended Data Figure 4.

Mutational signature assignment was performed for all SBS mutations assigned to phylogenetic trees treating each branch / edge as a unique sample, hence ensuring that mutations were not double counted. HDP mutational signature extraction with deconvolution into reference signatures was undertaken to define the proportion of each mutational signature in each sample. Deconvoluted signatures were used to define the relative probability of each trinucleotide context per signature. Mutational assignment probability was defined as the probability *P* that, that a particular mutation, *i*, could be assigned to a given signature, *j*, in genome *k* was calculated as follows:

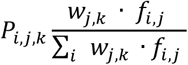

where *w*_*j,k*_ is the proportion of mutations assigned to signature *j* in genome *k* and *f*_*i,j*_ is the fraction of mutations in signature *j* that are the same substitution type and occur at the same trinucleotide context as mutation *i*.

### Cancer driver mutations

Cancer driver mutations were identified using two methods aiming to identify genes and mutations in this cohort that are subject to positive selection. Firstly, to identify mutations in cancer genes under positive selection in an unbiased manner, we ran a modified dNdS method^64^. To avoid double-counting of mutations, only unique mutations (SBS and ID) which were mapped to branches of the phylogenetic trees were analysed. dndscv was run using the following parameters; max_coding_muts_per_sample=5000 and max_muts_per_gene_per_sample=20. The mutational processes associated with defective DNA polymerases have a well-reported extended sequence context bias^8,64^ which alters the expected probability of observing a mutation in specific trinucleotide nucleotide contexts. To account for this bias, a modified dNdS method was applied. Global dNdS values for the expected number of each mutation type were replaced with corrected values taking into account the observed mutation subtype (synonymous, missense, nonsynonymous, and splice site) totals. A generalized negative binomial linear model was applied to each mutation subtype accounting for the biased distribution observed. P-values were combined using Fisher’s method and multiple testing correction was performed with Benjamini-Hochberg method. Genes with a qval of >0.05 were considered to be under positive selection.

A second phase of cancer gene mutation analysis was undertaken; identifying mutations in this cohort which are codified in cancer mutation databases and exhibit characteristic traits of cancer driver mutations; an approach previously employed in the study of normal tissues^28,29^. In this phase of the analysis we sought to identify the spectrum and frequency of cancer driver mutations in this cohort. Somatic mutations (SBS and ID) were collated per-sample from all tissues. Analysis was restricted to coding regions and mutations were filtered using lists of known cancer genes; mutations in samples from intestinal epithelium were filtered using a list of 90 genes associated with colorectal cancer^26^; samples from all other tissues including blood were filtered using a pan-cancer list of 369 driver genes^64^. Genes were then characterised according to their predominant molecular behaviour; dominant, recessive or intermediate (those demonstrating aspects of both types of behaviour) using the COSMIC Cancer Gene Census^65^. All candidate mutations were annotated using the cBioportal MutationMapper database (https://www.cbioportal.org/mutation_mapper). Mutations meeting the following criteria were considered to be driver mutations; truncating mutations (those that cause a shortened RNA transcript, nonsense, essential splice-site, splice region and frameshift ID) in recessively acting genes, known activating hotspot mutations in dominant (and recessive) genes and lastly mutations that were in neither of the above categories but characterised by the MutationMapper database as being ‘likely oncogenic’ were also included in the final driver mutation catalogue. We also sought to compare the frequency of driver mutations in histologically normal crypts with POLE and POLD1 mutations to those from individuals who do not carry DNA polymerase mutations. Somatic mutations from 445 normal intestinal crypts^26^ were annotated and filtered using the above criteria. Comparison was made with normal intestinal crypts from this cohort of individuals with POLE and POLD1 germline mutations (Extended Data Figure 6).

### Embryonic variant calling

Whole-genome sequencing of bulk blood samples were used to identify early embryonic SBS and ID mutations. Since bulk blood represents a very polyclonal tissue, variants found in blood reflect those generated in the first few cell divisions of life^52^. Variant counts from blood samples were included in the germline and artefact filtering as described above. For SBSs, a minimum VAF of 0.15 was required to be included in the embryonic set. Of the remaining SBSs, 205 out of a total 385 (53%) were shared with intestinal samples, confirming they must have arisen prior to gastrulation. For ID, we set the minimum to 0.1 to reflect the higher levels of noise accompanying indel calling and variant read counting. For indels, this amounted to 28 out of 30 (93%).

To investigate the role of POLE mutagenesis in the early embryo, we used the mutational signature contribution to the observed SBS counts, given the highly elevated SBS mutation rate. We fitted SBS1, SBS5, SBS10a, SBS10b and SBS28 to patient-specific embryonic counts using SigFit. SBS1 and SBS5 reflect the normal background mutagenesis already present in the embryo^52,66^, while the other signatures are caused by defective POLE.

For POLD1 mutagenesis, we quantified the number of insertions of T at homopolymers of T, the characteristic peak in ID1 and the one dominating the indel landscape in POLD1 patients. We used insertions rather than SBSs because of the relatively modest increase in SBS mutation rate, but a much higher increase in the rate of insertion acquisition.

### Telomere length estimation

Telomere attrition is a hallmark of cellular aging and is accelerated in certain disease processes. To assess the length of telomeres in the tissue samples in this cohort we undertook estimation of telomere lengths using two established methods of telomere length estimation from next-generation sequencing data.

Telomerecat is a ploidy-agnostic method of telomere length estimation (to base pair resolution) from next-generation sequencing data which has been benchmarked across human and animal studies in normal tissues and cancers^67^. This method has been employed in previous studies of somatic mutations in normal mutations^26-28^. We generated telomere length estimates for all samples using 100 simulator runs (parameter -N100). Results for most but not all samples were plausible and showed a positive correlation with those from a second telomere length content algorithm (TelomereHunter). Approximately 30% of samples returned zero values for telomere length, similar observations have been made in other data sets sequenced on the Illumina NovaSeq platform. Results of the algorithm based on sequencing data generated by the Illumina X10 and other sequencing platforms does not demonstrate this pattern and can be relied upon. For this analysis we favoured TelomereHunter which is well established method used in tumour sequencing analyses^68^, shows good concordance with other methods of telomere length estimation^69^ and is reliable across all tested samples sequenced on the Illumina NovaSeq platform.

Telomere content measurements were generated by running TelomereHunter using default parameters across all histologically normal crypts in this cohort (n=109) and normal crypts from a previous study that do not have DNA proofreading polymerase germline mutations (n=445)^26^. To assess age-related telomere attrition in normal tissues we fitted a linear mixed-effects model to assess the effect of age and also test whether telomere attrition is greater in crypts with a DNA proofreading polymerase mutation. Age was fitted as a fixed effect and patient as a random effect, an additional dichotomous genotype variable was added as a fixed effect. We assume a similar length at birth hence fitted a fixed intercept and assess the difference in slope between samples from this cohort and the non-predisposed crypts. We compare the model fit using ANOVA and the difference between models using a chi-squared test. P-value thresholds of >0.05 were used (Supplementary Code)

## CODE AVAILABILITY

Code required to reproduce the analyses in this paper are available online. Mutation calling algorithms are available through GitHub (https://github.com/cancerit). Bespoke R scripts described in this study are available online (https://github.com/TimCoorens/Polymerase). Code for statistical analyses is provided as part of the Supplementary code. All other code is available from the authors upon request.

## ACKNOWLEDGEMENT

We thank the staff of Wellcome Sanger Institute Sample Logistics, Genotyping, Pulldown, Sequencing and Informatics facilities for their contribution including Laura O’Neill, Yvette Hooks, Stephen Gamble, Calli Latimer and Kirsty Roberts for their support with sample management and laboratory work. In addition; Laura Humphreys and the Cancer Research UK (CRUK) Mutographs Grand Challenge team for their support with this study, Raheleh Rahbari for advice and guidance, Thomas Mitchell for advice regarding statistical analyses, Moritz Gerstung and Harald Vöhringer for help with analysis, advice and discussions. Kieren Allinson (Cambridge University Hospitals) for assistance with histopathological review. We thank Katherine Sherwood (Edinburgh Cancer Research Centre, IGMM, University of Edinburgh) and Laura Chegwidden (Institute of Cancer and Genomic Sciences, University of Birmingham) for their assistance in obtaining samples. We thank the participants of the CORGI and CORGI 2.0 studies and local investigators and their teams, without whose support this work would not be possible. We particularly thank Carole Brewer, Paul Lidder and Tamsin Pullen at Royal Cornwall Hospital Trust, Andrew Latchford, Huw Thomas and Ripple Man at St Marks Hospital, London, Maria Petmann and Angela Andrews at Nottingham, James East, Conor Lahiff and Hellen Purnell at Oxford University Hospitals and Jeanette Rothwell, Gareth Evans and James Hill at Manchester University NHS.

## CONTRIBUTIONS

P.S.R., T.H.H.C., M.R.S., I.T. and C.P. conceived the study design. C.P., Ly.M and I.T. recruited individuals, collected samples and curated sample and clinical data. P.S.R., B.L., J.H., C.M.A.P. and S.G undertook laboratory work. J.H. coordinated sequencing submissions and contributed to data management. L.M. developed bespoke DNA library methods. F.A. and I.M. developed bespoke sequencing methods and analysed data. L.M. and H.L.S. contributed and analysed control data. P.S.R., T.H.H.C., E.M., F.A., A.R.J.L., S.O. and M.A.S. performed data analysis. M.R.S., P.C., I.M. and M.A.S. oversaw statistical analysis. M.R.S. and I.T. oversaw the study. All authors were involved in the preparation and review of the manuscript.

## EXTENDED DATA FIGURES

**Extended Data Figure 1.**
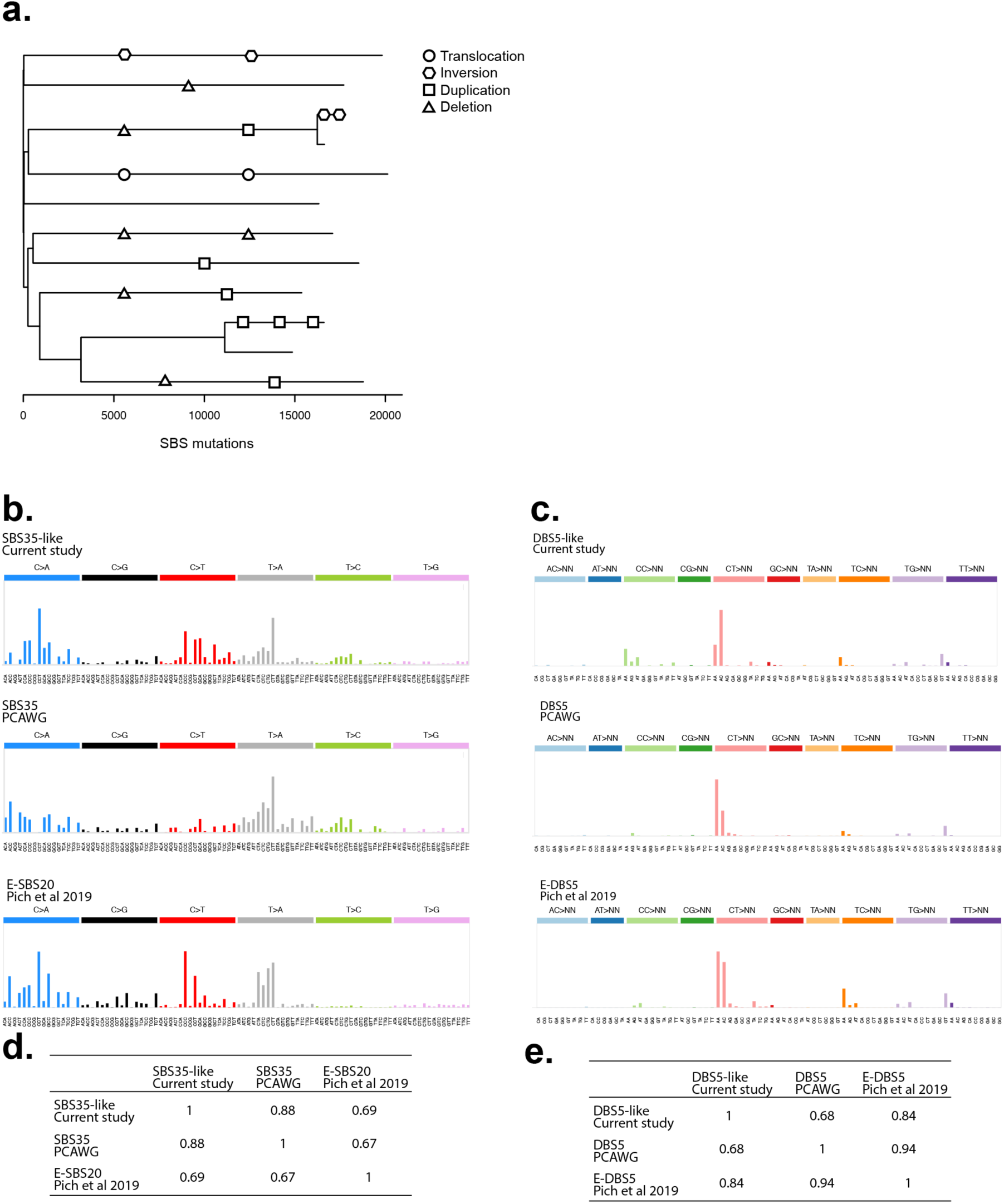
Mutational signatures and structural variants in normal tissue from an individual exposed to oxaliplatin chemotherapy. Mutational signatures identified in normal intestinal crypts from individual PD44592 who had previously undergone treatment with oxaliplatin. Distinctive SBS and DBS signatures were observed that have previously been associated with oxaliplatin treatment. Unexpectedly high numbers of structural variants were also observed in the normal intestinal crypts from this individual and may be due to chemotherapy exposure. **(a)** Phylogenetic tree from individual PD44592 with structural variants (SVs) mapped to branches. Shapes correspond to SV type; translocation (circle), inversion (hexagon), large duplication (square) and deletion (triangle). Ordering of SVs on branches is arbitrary **(b)** SBS signature SBS35-like displayed above the Pan-Cancer Analysis of Whole Genomes (PCAWG) reference signature SBS35 (PCAWG)^9^ and E-SBS20^43^ **(c)** Doublet Base Substitution (DBS) signature displayed above PCAWG reference signature DBS5 and E-DBS5 from Pich et al 2019^43^. **(d and e)** Cosine similarity matrices for SBS and DBS signatures.

**Extended Data Figure 2.**
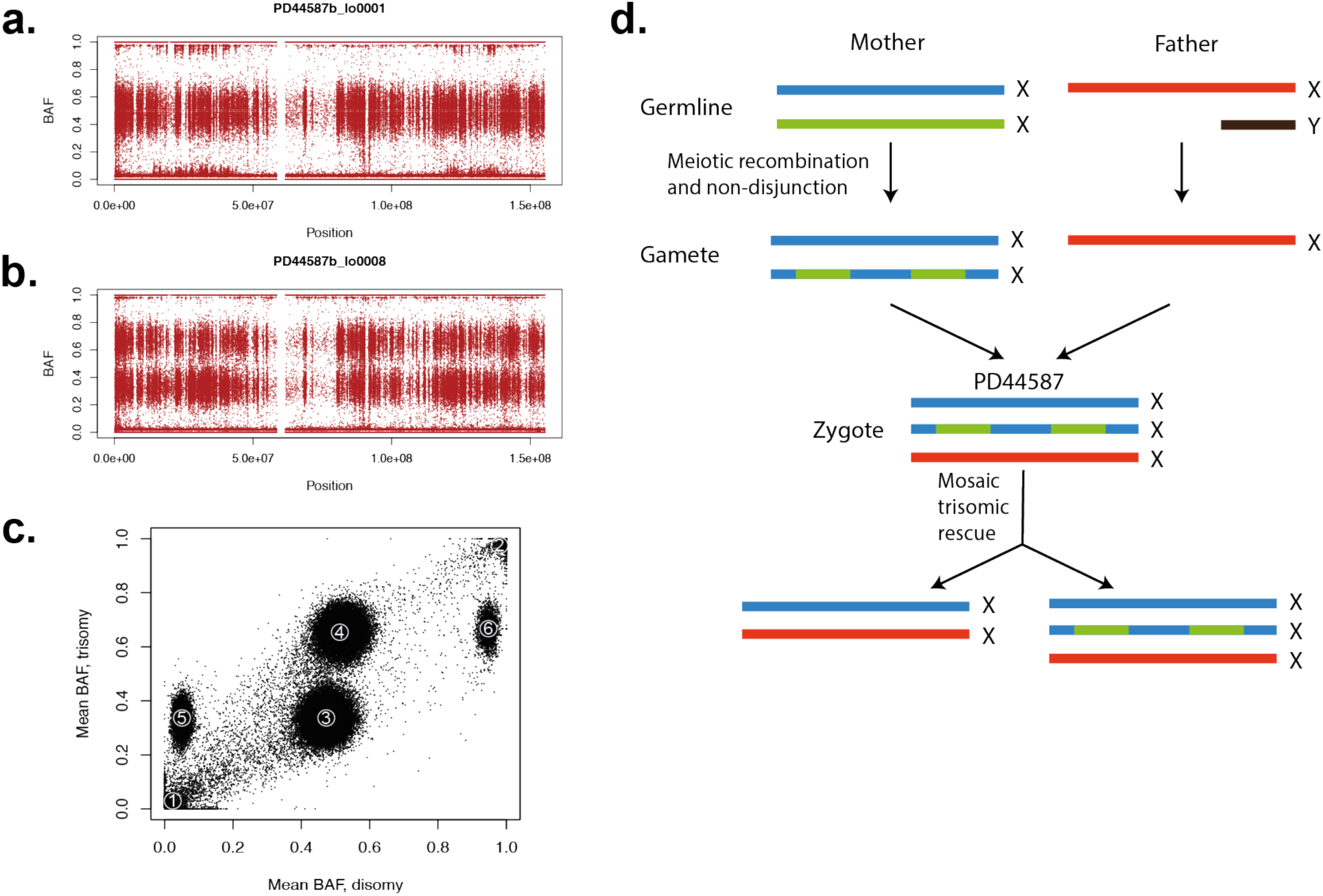
Trisomy X (47XXX) with mosaic trisomic rescue in individual PD44587 identified by lineage tracing of somatic mutations. B-allele frequencies (BAF) of SNP sites for intestinal crypts with two copies of the X-chromosome **(a)** or three **(b)**. Seven crypts exhibited the disomy, whereas one crypt and the majority of blood showed the trisomy. **(c)** The mean BAF of SNPs for samples with a disomic profile versus those with a trisomic profile. SNPs clustered in six distinct groups: those absent from all samples (marked with ‘1’), homozygously present across all samples (2), heterozygous in disomy but in one out of three copies in trisomy (3), heterozygous in disomy but on two copies in trisomy (4), absent from disomy but on one copy in trisomy (5) or homozygous in disomy but on two copies in trisomy (6). The last two clusters are inconsistent with an acquired gain of chromosome X, as they constitute bringing in novel germline SNPs or omission of those previously homozygotic. **(d)** Therefore, this profile can only be explained by a zygote which possessed three copies of X, one of which was mosaically lost in the crypt lineage.

**Extended Data Figure 3.**
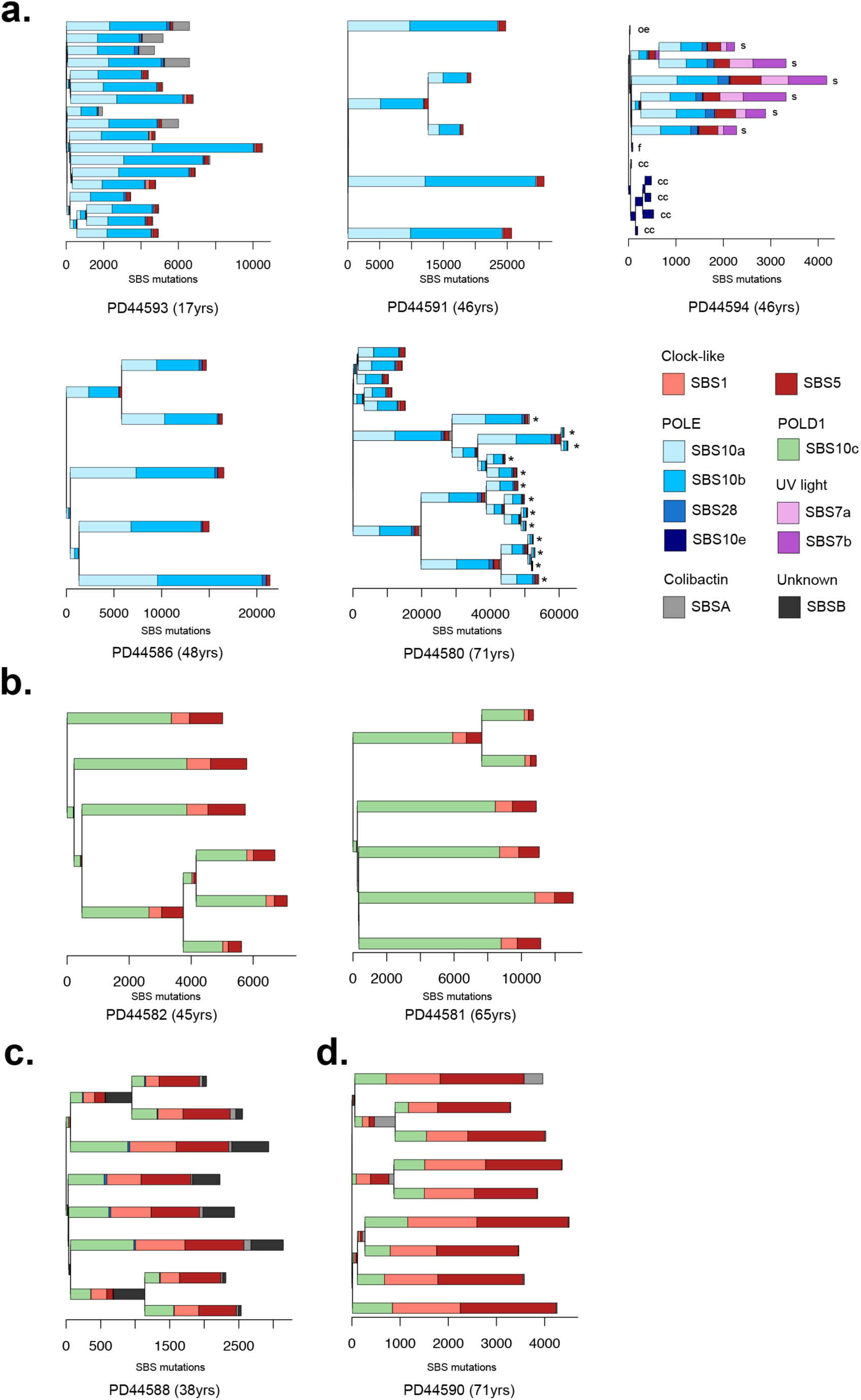
Phylogenetic trees constructed from somatic single base substitution (SBS) mutations annotated with mutational signature exposure. Phylogenetic trees generated from normal and neoplastic crypts displayed per individual. Mutational signature exposure per branch is displayed as stacked bar plots. Ordering of mutational signatures within branches is arbitrary. SBS burden is displayed on the x-axis. Each branch tips represent normal intestinal crypts unless otherwise indicated microbiopsies from; ‘s’ skin, ‘cc’ cerebral cortex, ‘oe’ oesophagus, ‘f’ hair follicle and ‘*’ indicates adenomatous crypts. Trees are grouped according to germline DNA polymerase mutation; **(a)** POLE L424V, **(b)** POLD1 S478N, **(c)** POLD1 L474P and **(d)** POLD1 D316N.

**Extended Data Figure 4.**
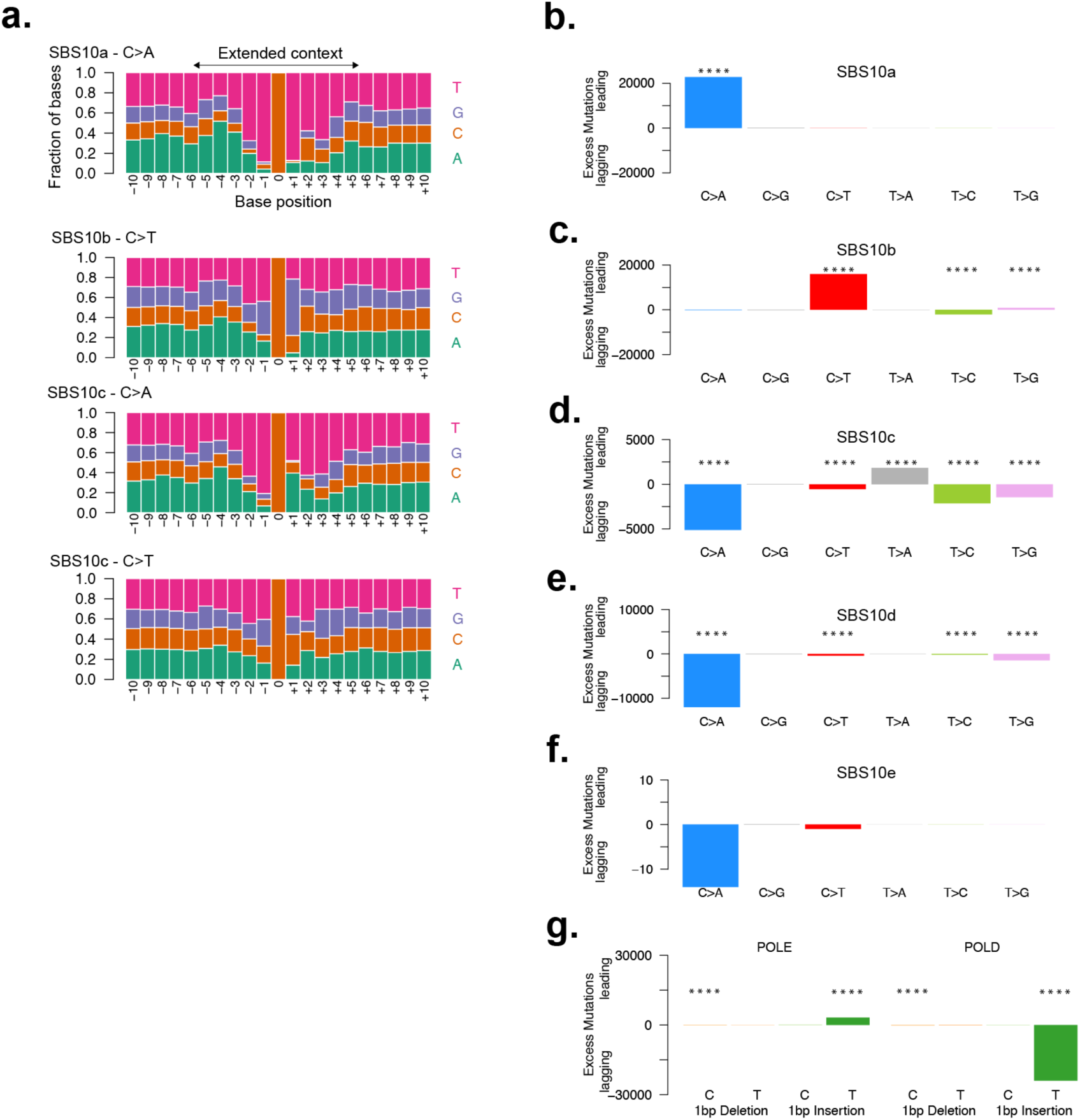
Characterisation of replication strand bias and extended sequence context of mutational signatures associated with germline DNA polymerase mutations. **(a)** Extended sequence context of mutations assigned to signatures SBS10a, 10b and SBS10c displayed with pyrimidine annotation. Plots show extended sequence context of C>A and C>T mutations in SBS10a and SBS10b respectively and both C>A and C>T mutations in signature SBS10c. **(b-f)** Replication strand biases of SBS mutations indicated by excess mutations on either the leading or lagging DNA strand. Biases are displayed according to mutations assigned to known signatures SBS10a and SBS10b and new signatures SBS10c, SBS10d and SBS10e. Only SBS mutations with an assignment probability >0.7 were included (Methods). Mutations are classified according to their pyrimidine annotation. Strong replicative strand asymmetries are seen in known (SBS10a & SBS10b) signatures as well as new signatures; SBS10c and SBS10d. Mutation classes with statistically significant replication strand bias (adjusted p < 0.0001) are annotated with ‘****’. P-value correction for multiple testing was performed using the Benjamini-Hochberg method. No significant replicative strand asymmetry is observed in SBS10e. **(g)** ID replication strand asymmetries are displayed for single base insertions and deletions according to the affected polymerase gene; POLE (left) POLD1(right). Replication strand bias is indicated by excess mutations with single base insertions on the leading strand in POLE mutant cells and on the lagging strand in POLD1 mutant cells.

**Extended Data Figure 5.**
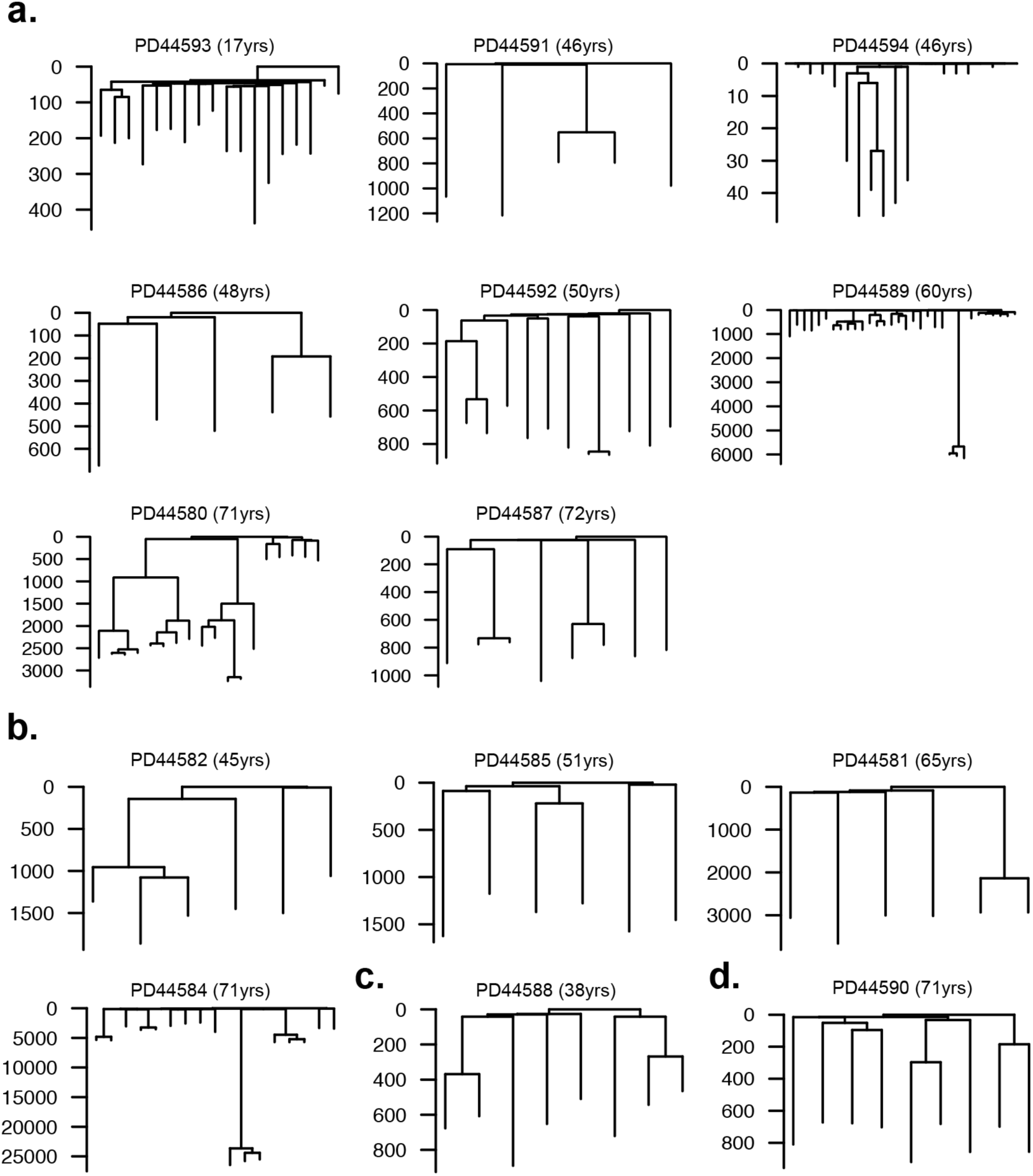
Phylogenetic trees constructed from somatic insertion and deletion (ID) mutations from normal and neoplastic intestinal stem cells. Phylogenetic trees generated from ID mutations. ID mutation burden is displayed on the y-axis. Each tree represents samples from a single individual. Trees are grouped according to germline DNA polymerase mutation **(a)** POLE L424V **(b)** POLD1 S478N **(c)** POLD1 L474P **(d)** POLD1 D316N.

**Extended Data Figure 6.**
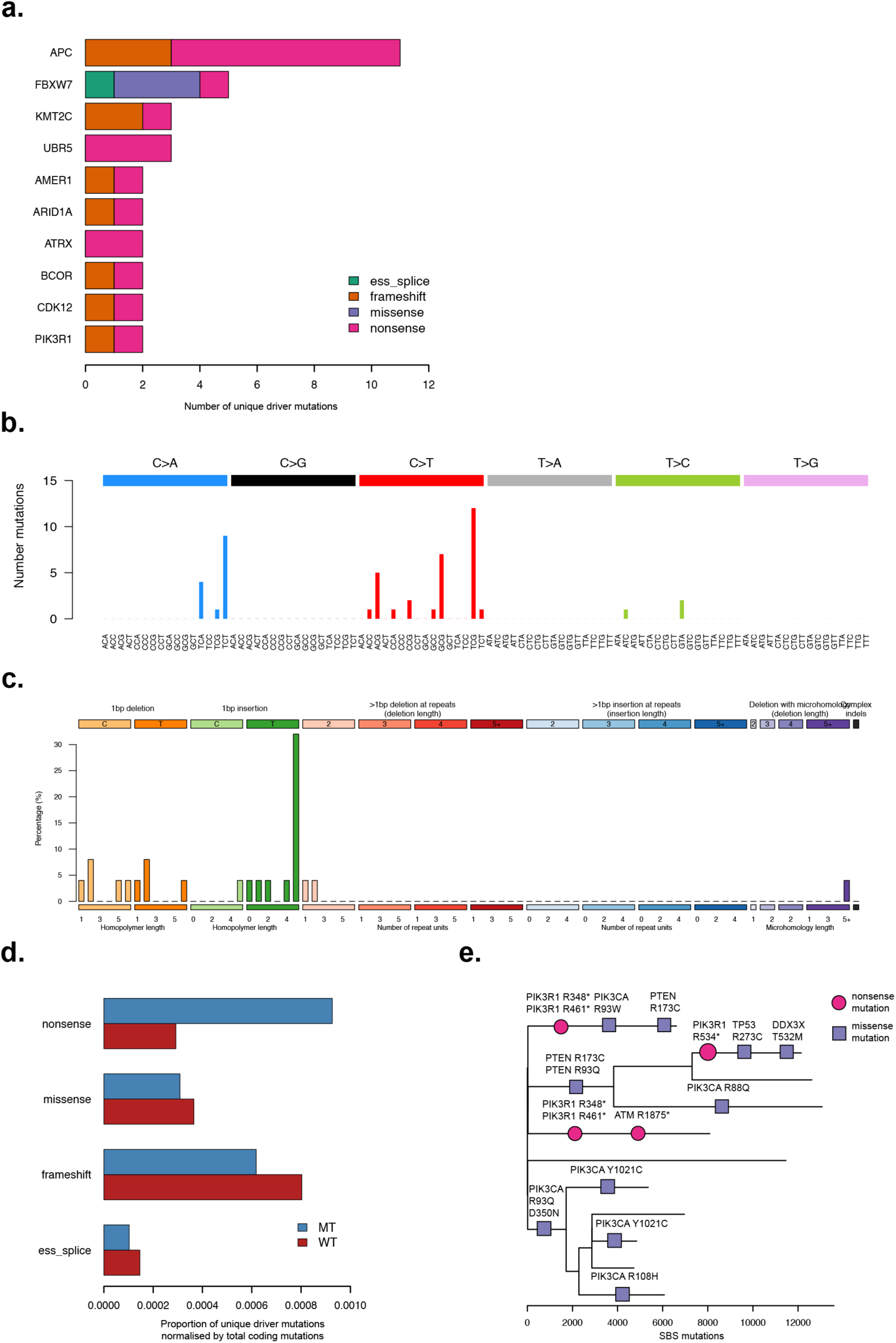
Driver mutation landscape in normal tissues from individuals with POLE and POLD1 germline mutations. Driver mutations in tissues from individuals with germline DNA polymerase mutations. **(a)** Cancer driver mutations (top 10) identified in histologically normal intestinal crypts displayed in order of frequency, mutation class is indicated by the colour. **(b)** Trinucleotide mutational spectrum of SBS driver mutations from normal and neoplastic cells showing characteristic peaks associated with DNA polymerase mutational signatures. **(c)** ID mutation spectrum showing ID type of driver mutations are associated with ID1 mutational signature. **(d)** Comparison of driver mutation burden of normal crypts from individuals with a germline DNA polymerase mutation (‘MT’, blue) vs. wild-type crypts (‘WT’, red). Proportion of driver mutations is displayed on the x-axis normalised for the total burden of coding SBS mutations. SBS mutations included are from n=445 wild-type intestinal crypts^26^ and n=109 and DNA polymerase mutant crypts from the current cohort. **(e)** Phylogenetic tree of SBS mutations from endometrial glands from individual PD44589. SBS driver mutations are plotted on the tree, ordering of the drivers within each branch is arbitrary. Driver mutation class is represented by the symbol: nonsense mutations (circles) missense (squares). Gene name and protein change are displayed above each symbol.

**Extended Data Figure 7.**
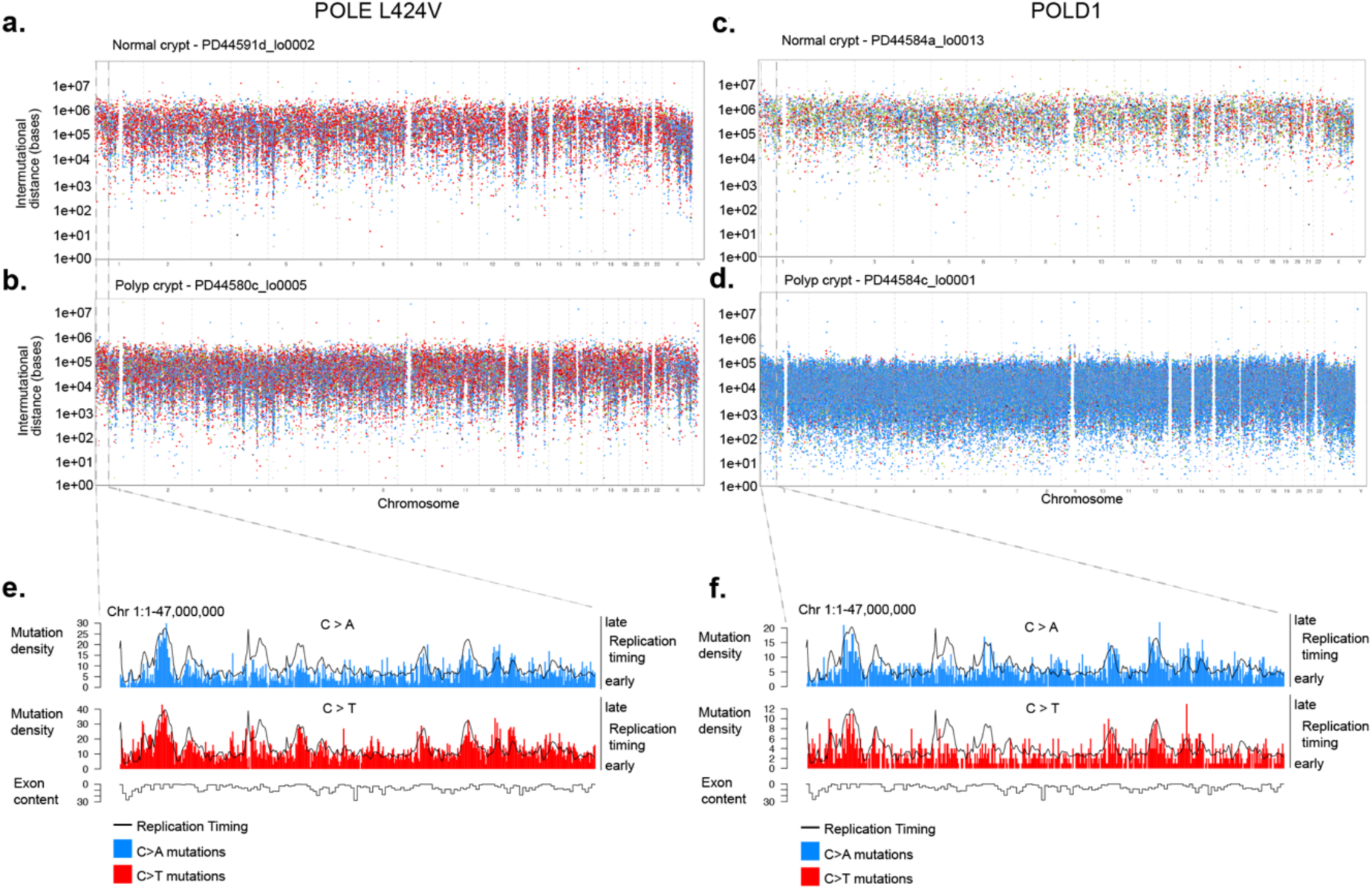
Genome distribution of mutational processes associated with germline DNA polymerase mutations. **(a-d)** Rainfall plots showing inter-mutational distance of SBS mutations plotted on a logarithmic scale per chromosome (on x-axis). Dots are coloured according to SBS mutation type; C>A (blue),C>G (black),C>T (red),T>A (grey), T>C (green), T>G (pink). Plots are displayed per genome for four representative crypts according to their germline mutation and histological type. **(e-f)** Bar and line plots of mutation density, replication timing and exon density. Region displayed is from chromosome 1 and shows increased mutation density correlating with late replicating regions and reduced exon density. Mutation density is displayed in 10kb bins, exon content (percentage of bin) is displayed across a 3-bin width. Mutation data is aggregated according to mutated polymerase gene and displayed according to mutation type; C>A (blue) and C>T (red). DNA Replication timing of each bin is demonstrated by a single black line (right-hand axis).

**Extended Data Table 1** | **Clinical summary and phenotypic characteristics of individuals with germline POLE and POLD1 EDM mutations**

Summary of phenotypic features and disease burden in all individuals in this cohort as well as those published to date.

**Extended Data Table 2** | **Summary of samples including germline DNA polymerase mutation, sequencing method and mutation burdens**

**Extended Data Table 3** | **Cancer driver mutations identified in this cohort**

Cancer driver mutations identified across all cell types studied in this cohort.

